# Targeting BRPF3 moderately reverses olaparib resistance in high grade serous ovarian carcinoma

**DOI:** 10.1101/2021.12.21.473688

**Authors:** Benjamin G. Bitler, Tomomi M. Yamamoto, Alexandra McMellen, Hyunmin Kim, Zachary L. Watson

## Abstract

**Background:** PARP inhibitors (PARPi) kill cancer cells by stalling DNA replication and preventing DNA repair, resulting in a critical accumulation of DNA damage. Resistance to PARPi is a growing clinical problem in the treatment of high grade serous ovarian carcinoma (HGSOC). Acetylation of histone H3 lysine 14 (H3K14ac) and associated histone acetyltransferases (HATs) and epigenetic readers have known functions in DNA repair and replication. Our objectives are to examine their expression and activities in the context of PARPi-resistant HGSOC, and to determine if targeting H3K14ac or associated proteins has therapeutic potential.

**Results:** Using mass spectrometry profiling of histone modifications, we observed increased H3K14ac enrichment in PARPi-resistant HGSOC cells relative to isogenic PARPi-sensitive lines. By RT-qPCR and RNA-Seq, we also observed altered expression of numerous HATs in PARPi-resistant HGSOC cells and a PARPi-resistant PDX model. Knockdown of HATs only modestly altered PARPi response, although knockdown and inhibition of PCAF significantly increased resistance. Pharmacologic inhibition of HBO1 severely depleted H3K14ac but did not affect PARPi response. However, knockdown and inhibition of BRPF3, which is known to interact in a complex with HBO1, did reduce PARPi resistance.

**Conclusions:** This study demonstrates that severe depletion of H3K14ac does not affect PARPi response in HGSOC. Our data suggest that bromodomain functions of HAT proteins, such as PCAF, or accessory proteins, such as BRPF3, may play a more direct role compared to direct histone acetyltransferase functions in PARPi response.

## Background

High grade serous ovarian carcinoma (HGSOC) is the deadliest gynecological malignancy. It is the fifth most common cause of cancer death in women and has one of the highest death-to-incidence ratios of all cancers (1). Most patients respond initially to standard practice cytoreductive surgery and platinum/taxol chemotherapy (2). The addition of PARP inhibitors (PARPi) as first-line maintenance therapy, supported by numerous clinical trials in homologous recombination-deficient and -proficient cohorts, has extended progression-free survival intervals for many patients (3–5). Despite these advances, a substantial proportion of patients do not respond to PARPi, and over 80% of cases recur and are expected to acquire resistance to all available therapies (2, 6). Since nearly all HGSOC patients will be eligible to receive PARPi, it is essential to discover mechanisms of resistance and provide support for novel therapies to overcome resistance and improve outcomes.

Numerous studies and trials have focused on exploiting vulnerabilities to augment or restore PARPi sensitivity, including the use of topoisomerase inhibitors (7, 8), cyclin-dependent kinase knockdown or inhibition (9, 10), Wnt inhibitors (11), and histone methyltransferase inhibitors (12), among others [reviewed in (6)]. In the current study we examine histone modifications and expression of histone modifying enzymes in the context of PARPi-resistant HGSOC. Histone modifications play critical roles in PARPi resistance, as they control access to genomic DNA for vital processes including transcription, replication, and repair (13, 14). When dysregulated, these same modifications have been shown to contribute to every aspect of cancer biology, from early oncogenesis to progression, metastasis, chemoresistance, and immune evasion (15–18). Acetylation of histone H3 lysine 14 (H3K14ac) neutralizes the positively charged lysine residue and reduces interactions between the histone octamer and encircling DNA, thus allowing greater access for transcriptional, replicative, and DNA repair machinery (19, 20). As such, H3K14ac is generally a modification associated with transcriptionally active chromatin.

While H3K14ac is only a single modification, its regulation is complex, as it may be catalyzed by numerous histone acetyltransferases (HATs) of different classes including (but not limited to) GCN5 and PCAF (encoded by *KAT2A* and *KAT2B*, respectively), CBP and P300 (encoded by *KAT3A* and *KAT3B*, respectively), as well as HBO1, also known as MYST2 (encoded by *KAT7*). H3K14ac is known to be a key factor in the process of single-stranded DNA break (SSB) and double-stranded DNA break (DSB) repair. For SSB repair, H3K14ac stabilizes chromatin interactions and promotes error-free Nucleotide Excision Repair (NER) (21). HATs GCN5 and PCAF are structurally similar and promote repair by acetylation of H3K14, and also by acetylation of non-histone targets including Replication Protein A1 (RPA1) (22). Acetylation of RPA1 by GCN5/PCAF activates NER and promotes accumulation of DNA repair proteins at sites of damage. For DSB, CBP and P300 are structurally similar and promote chromatin remodeling and recruitment of effectors involved in both homologous recombination (HR) and non-homologous end joining (NHEJ) (23, 24). Additionally, CBP/P300 promote transcription of HR factors BRCA1 and RAD51 recombinase (25). Finally, phosphorylated HBO1 has been shown to facilitate recruitment of the NER protein XPC to sites of cyclopyrimidine dimers (CPD) and is essential for NER at sites of ultraviolet (UV) irradiation damage (26).

Modifiers of histone acetylation, especially histone deacetylases (HDACs) and HDAC inhibitors, have been investigated as therapeutic targets in HGSOC [reviewed in (27)]. However, the role of H3K14ac has not been studied in the context of PARPi resistance. Our objective is to examine alterations to H3K14ac, including changes in expression or activity of associated epigenetic writers and readers, and to determine the therapeutic potential of perturbing H3K14ac in PARPi-resistant HGSOC.

## Results

### H3K14ac enrichment and expression of HATs are altered in PARPi-resistant HGSOC cell lines and PDX models

Three *TP53*-mutated HGSOC cell lines (PEO1, OVSAHO, and OVCA433) were subjected to stepwise dose escalation of the PARPi olaparib to select a population of resistant cells. PEO1 cells encode *BRCA2* nonsense mutation 5193C>G and are BRCA2-deficient (28); OVSAHO have a *BRCA2* homozygous deletion (29); OVCA433 are positive for BRCA1 and BRCA2. Olaparib resistance was confirmed in each line using dose response colony formation assays. Olaparib response in PEO1 (IC_50_ = 6.4 nM) and PEO1-OR (IC_50_ = 1223 nM), and mechanisms of resistance have been reported (11, 12). The olaparib IC_50_ increased from 322 nM in OVSAHO to 1300 nM in OVSAHO-OR, and from 129 nM in OVCA433 to 5580 nM in OVCA433-OR. Representative olaparib dose response curves and calculated IC_50_ values are shown in **Supplemental Fig. S1**. Histones from each parental and olaparib-resistant line were isolated and the percentage of histone H3 and H4 modifications were determined by mass spectrometry. We reported these data for PEO1/PEO1-OR (12), and we include these data here for comparison with the two new sensitive/resistant isogenic pairs. Complete mass spectrometry data for PEO1/PEO1-OR are published (12), and data for all marks analyzed in the new lines are available in **Supplemental File 1**. All samples were analyzed in triplicate, yielding a “Total Average Percent” value. This number represents the percentage of the specific modification divided by the total detection of a histone tail residue. For example, in OVSAHO cells, H3K14ac was detected as 19.2% of all H3K14 **(Fig. 1A** and **Supplemental File 1)**. Notably, H3K14ac was significantly increased in all three resistant lines compared to the parental lines **(Fig. 1A)**. We subsequently isolated histones from each cell line pair and immunoblotted for H3K14ac and total H3. By performing sensitive densitometry to detect band intensity and normalizing to total H3, we confirmed increased H3K14ac in all three olaparib-resistant lines **(Fig. 1B)**.

**FIG 1.**
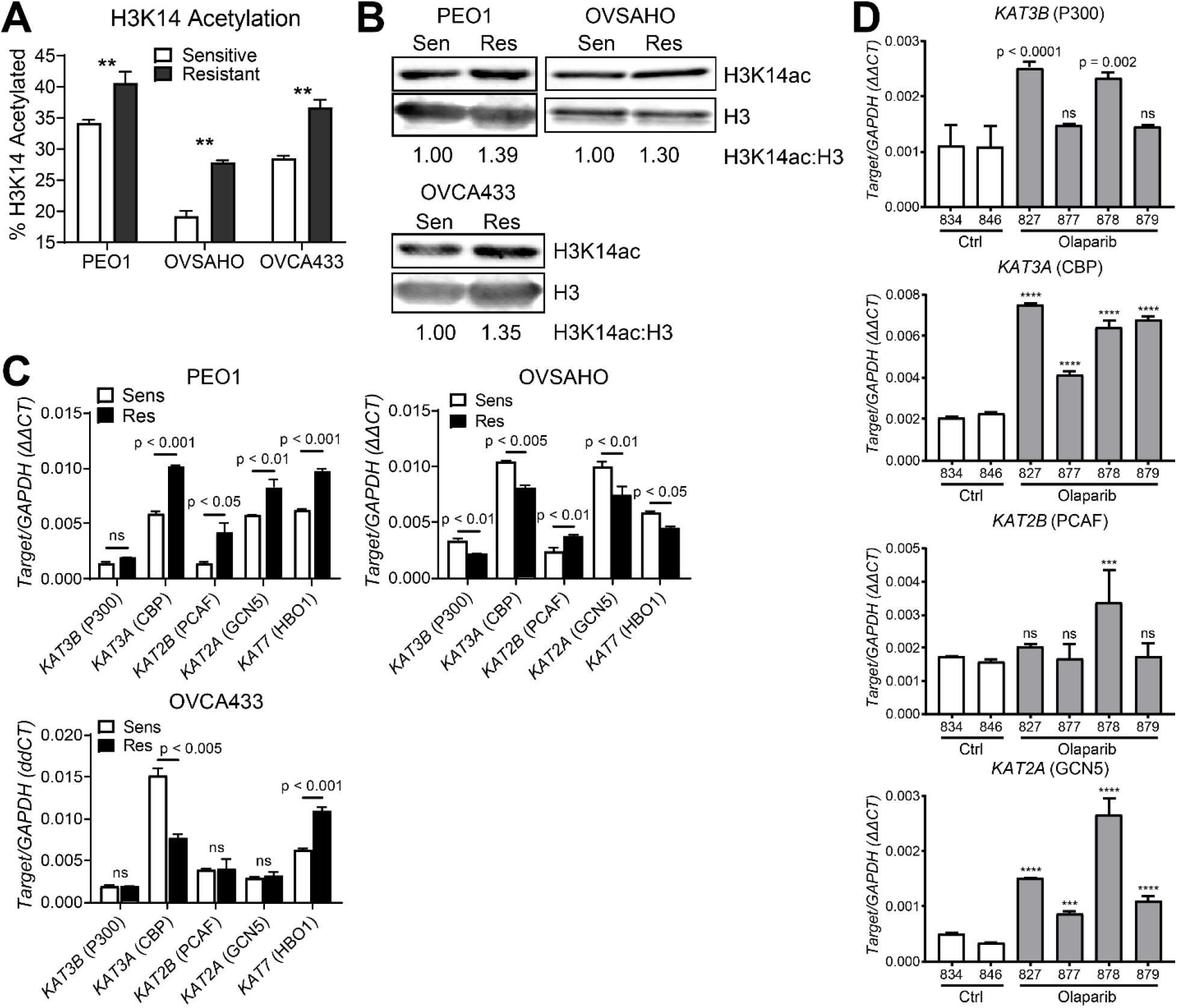
H3K14ac enrichment and histone acetyltransferase expression is altered in olaparib-resistant HGSOC cells and a PDX model. **(A)** Histone extracts from the indicated olaparib-sensitive and olaparib-resistant HGSOC cell lines were assayed in triplicate by mass spectrometry for histone modifications. Data are shown for each sensitive/resistant pair as the mean ± SD percentage of acetylated H3K14. N = 3; p-value by t-test. ** p < 0.01. **(B)** Histone extracts from the indicated olaparib sensitive/resistant pairs were assayed by Western blotting for H3K14ac and total H3. The H3K14ac:H3 ratio was determined by band densitometry and normalized to the olaparib-sensitive line of each pair, which are set as 1. **(C)** mRNA from the indicated olaparib-sensitive/-resistant pairs was assayed for HAT expression by RT-qPCR. Target gene expression was determined relative to *GAPDH* control by ΔΔCt method. Each bar represents the mean ± SD of three PCR reactions. p-values by t-test. **(D)** Human HGSOC PDX cells were injected into NSG mice. After seven days, mice received a 21-day treatment course of intraperitoneal vehicle control or 50 mg/kg olaparib. Following treatment, tumor cells were allowed to recur over an additional 21 days. Upon necropsy, ascites cells were collected. mRNA from ascites cells was analyzed by RT-qPCR for the indicated HATs. Target gene expression was determined relative to *GAPDH* control by ΔΔCt method. Each bar represents the mean ± SD of three PCR reactions. Numbers below bars are mouse ear tag IDs. p-values were determined by t-test comparing each olaparib-treated mouse (gray bars; #827, #877, #878, or #879) to the average of both control mice (white bars; #834 and #846). *** p < 0.001; **** p < 0.0001.

We next examined expression of HATs in each sensitive/resistant cell line pair. We examined expression of five HATs with known H3K14 acetylation activity by RT-qPCR, including *KAT3B* (P300), *KAT3A* (CBP), *KAT2B* (PCAF), *KAT2A* (GCN5), and *KAT7* (HBO1) (30–35). We found significantly different expression of at least two HATs in each cell line **(Fig. 1C)**. For example, PEO1-OR showed upregulation of all HATs except *KAT3B* relative to the olaparib-sensitive PEO1 cells. None of the HATs were predictive of up- or down-regulation of H3K14ac in the resistant cell lines. For example, *KAT7* was upregulated in PEO1-OR and OVCA433-OR but downregulated in OVSAHO-OR.

To establish an olaparib-resistant animal model, we inoculated NOD SCID gamma (NSG) mice by intraperitoneal injection with GTFB-PDX1009, a *TP53*-mutated, BRCA-wildtype HGSOC. We then treated tumor bearing mice daily with olaparib or vehicle control for 21 days, then monitored the mice for tumor recurrence and growth for two months [schematic published in (12)]. After recurrence and necropsy, tumor cells isolated from olaparib-treated mice were subsequently shown to be highly olaparib-resistant (11). By RT-qPCR, we found that mRNA expression of *KAT3A* and *KAT2A* were increased in all four olaparib-treated ascites, while *KAT2B* and *KAT3B* were upregulated in one and two mice, respectively **(Fig. 1D)**. We further performed RNA-Seq to compare the transcriptomes of control mice (#834 and #836) with two olaparib-treated mice (#827 and #878). RNA-Seq data confirmed significant upregulation of *KAT2A* (GCN5) and *KAT3A* (CBP) in the olaparib-treated mice **(Supplemental Table S1)**. In summary, H3K14 acetylation and HAT expression are altered in PARPi-resistant models of HGSOC, including BRCA-deficient and -proficient cell lines, as well as a BRCA-proficient PDX.

### Genetic knockdown of HATs in olaparib-resistant HGSOC cells moderately alters olaparib response

H3K14ac is associated with multiple DNA repair mechanisms. Augmented DNA repair is an important component of PARPi resistance (6), and we have previously shown that knockdown or inhibition of epigenetic enzymes such as histone methyltransferases can significantly improve olaparib response (12). We therefore chose to knock down each HAT highlighted in **Fig. 1C** and examine olaparib response. Since PEO1-OR showed upregulation of four of the five HATs, we used lentiviral transduction to stably express short hairpin RNAs (shRNA) targeting each HAT, or a non-targeting scrambled shRNA as control (shCTRL), in these cells. We used two independent shRNA constructs per HAT; The RNAi Consortium Numbers and target sequences are given in **Supplemental Table S2**. We determined the degree of knockdown for each shRNA relative to shCTRL by RT-qPCR **(Fig. 2A-E)**. We then performed dose response colony formation assays. Cells were dosed with olaparib ranging from 4.8 nM to 30 μM or vehicle control and the olaparib IC_50_ was calculated for each shRNA. Representative olaparib IC_50_ values are shown in **Table 1** and full olaparib dose response curves are shown in **Supplemental Figure S2A**. Both shRNA against *KAT3B* (P300), one shRNA against *KAT3A* (CBP), and both shRNA against *KAT2A* (GCN5) showed a statistically significant (p < 0.05) reduction in olaparib IC_50_ relative to shCTRL. However, sensitization in such significant cases was modest, with IC_50_ values remaining at 42% to 65% of shCTRL. We performed a second set of dose response experiments to determine colony formation specifically at 120 nM, 600 nM, and 3 μM olaparib **(Supplemental Figure S2B)**. Colony formation for all shRNA was not different from shCTRL at 120 nM or 600 nM. sh*KAT3A*#1 and sh*KAT2A*#1 showed small but significant reductions in PEO1-OR colony formation in 3 μM olaparib.

**FIG 2.**
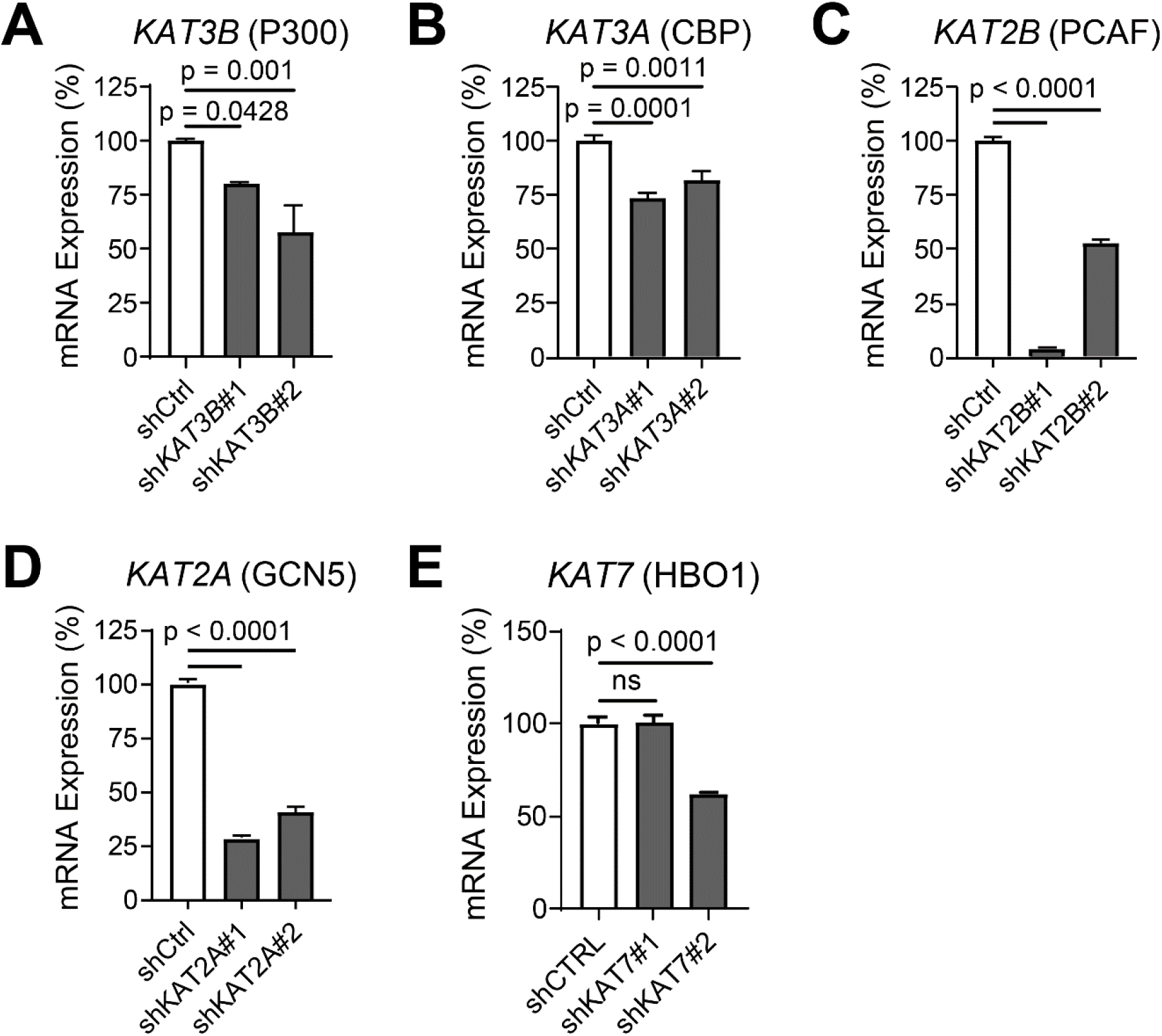
Knockdown of HATs shows only moderate alteration of olaparib response in PEO1-OR (olaparib-resistant) HGSOC cells. **(A-E)** PEO1-OR cells were transduced with the indicated targeted shRNA or a non-targeting scrambled shRNA (shCTRL) and then selected in puromycin. Knockdown of each HAT was determined by RT-qPCR. mRNA expression was determined relative to *GAPDH* by ΔΔCt method, then normalized to shCTRL for each knockdown. Each bar represents the mean ± SD of three PCR reactions. p-value by t-test.

**Table 1.** Olaparib IC_50_ values of control and HAT knockdown PEO1-OR HGSOC cells. For each indicated shRNA, the olaparib IC_50_ values are shown. Also shown are IC_50_ percentage relative to shCTRL (% shCTRL) and p-value. The column “2^nd^ Run?” indicates if the shRNA significantly changed colony formation in a follow-up experiment.

| **shRNA** | **Olaparib IC<sub>50</sub> (nM)** | **% shCTRL** | **p-value** | **2<sup>nd</sup> Run?** |
| --- | --- | --- | --- | --- |
| shCTRL | 2620 | - | - | - |
| shKAT3B #1 | 1699 | 65% | 0.016 | No |
| shKAT3B #2 | 1716 | 65% | 0.021 | No |
| shKAT3A #1 | 1672 | 64% | 0.014 | Yes |
| shKAT3A #2 | 1955 | 75% | 0.099 | No |
| shKAT2B #1 | 4549 | 174% | 0.002 | Yes |
| shKAT2B #2 | 3409 | 130% | 0.105 | Yes |
| shKAT2A #1 | 1104 | 42% | < 0.0001 | Yes |
| shKAT2A #2 | 1704 | 65% | 0.009 | No |
| **shRNA** | **Olaparib IC<sub>50</sub> (nM)** | **% shCTRL** | **p-value** |  |
| shCTRL | 1593 | - | - | - |
| shKAT7 #1 | 1431 | 90% | 0.444 | No |
| shKAT7 #2 | 1348 | 85% | 0.352 | No |

Despite differing levels of knockdown, both shRNA against *KAT2B* (PCAF) increased olaparib IC_50_ in PEO1-OR cells, with *shKAT2B*#1 showing a significant increase of 174% relative to shCTRL **(Table 1** and **Supplemental Figure S2A)**. This result was confirmed for both *KAT2B* shRNA in PEO1-OR in the follow up experiment, as both showed increased colony formation in 3 μM olaparib **(Supplemental Figure S2B)**. Neither shRNA targeting *KAT7* (HBO1) showed a significant effect in either experiment. In summary, we determined that specific knockdown of CBP (using sh*KAT3A*#1) and GCN5 (using sh*KAT2A*#1) modestly improved olaparib response, while knockdown of PCAF increased the olaparib IC_50_. All other knockdowns did not significantly alter olaparib response in PEO1-OR.

### Depletion of H3K14ac does not sensitize olaparib-resistant HGSOC cells, but bromodomains may play roles in olaparib response

We next determined if depletion of H3K14ac was sufficient to sensitize PEO1-OR cells to olaparib. We determined that 10 μM HBO1 inhibitor WM-3835, characterized by MacPherson et al. (36), severely depleted H3K14ac. By immunoblot densitometry, only 27% of H3K14ac remained in PEO1-OR cells after a 6-hour incubation **(Fig. 3A)**. We then performed colony formation assays using olaparib alone or in combination with 10 μM WM-3835. Despite the efficacy of H3K14ac depletion, no change in olaparib IC_50_ was found between the two treatments **(Fig. 3B and Table 2)**. Rather than continuing with direct inhibition of acetyltransferase activity, we elected to test if pharmacologic inhibition of the bromodomains (BRD) of HAT enzymes impacted olaparib response. We performed colony formation assays in PEO1-OR using olaparib alone or in combination with 2 μM GCN5/PCAF BRD inhibitor GSK4027, or 2 μM P300/CBP BRD inhibitor SGC-CBP30. GSK4027 notably increased olaparib resistance in PEO1-OR cells **(Fig. 3C and Table 2)**, while SGC-CBP30 had no effect on olaparib IC_50_ **(Fig. 3D and Table 2)**. This agrees with our data showing that knockdown of PCAF also increases olaparib resistance in these cells.

**FIG 3.**
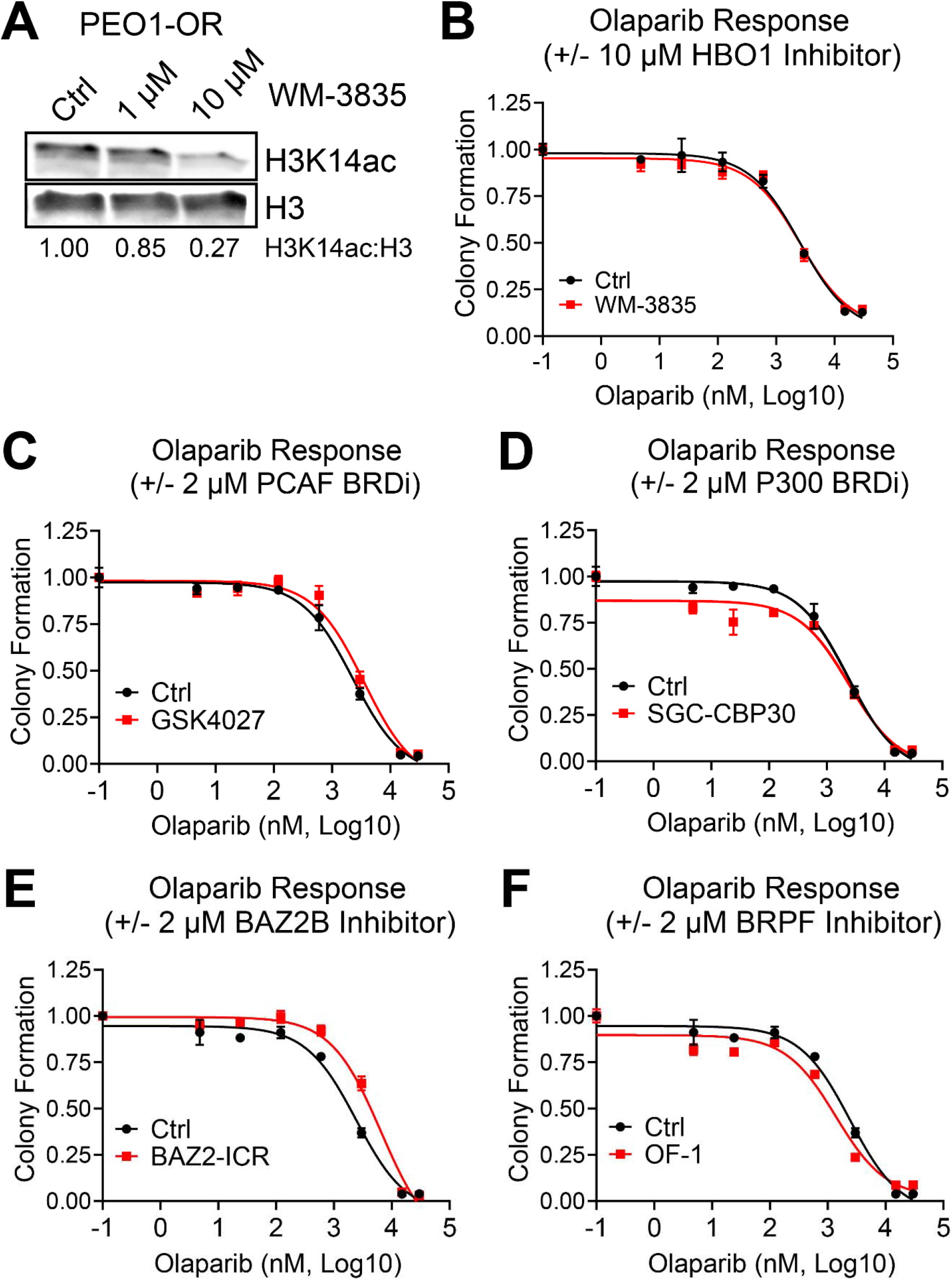
Severe depletion of H3K14ac does not shift olaparib IC_50_, but specific BRD inhibitors moderately affect olaparib response in PEO1-OR HGSOC cells. **(A)** PEO1-OR cells were treated with the indicated doses of HBO1 acetyltransferase inhibitor WM-3835 for 6 hours. Histone extracts were analyzed by Western blot for H3K14ac and total H3. The H3K14ac:H3 ratio was calculated by densitometry to quantify H3K14ac depletion. **(B)** PEO1-OR were seeded at 2500 cells per well in a 24-well plate and treated with increasing doses of olaparib with or without 10 μM WM-3835. Media and drug were changed every 2-3 days for eight days, after which colonies were fixed and stained with crystal violet. Stain was dissolved and the absorbance from each well was read by spectrophotometer, then normalized to vehicle control. Error bars, SEM. Olaparib IC_50_ for each treatment condition was calculated in GraphPad Prism. **(C-F)** PEO1-OR cells in a 24-well plate were treated with increasing doses of olaparib with or without 2 μM of the indicated BRD inhibitors. Colony formation and olaparib IC_50_ were determined as described for Fig. 3B. Error bars, SEM.

**Table 2.**
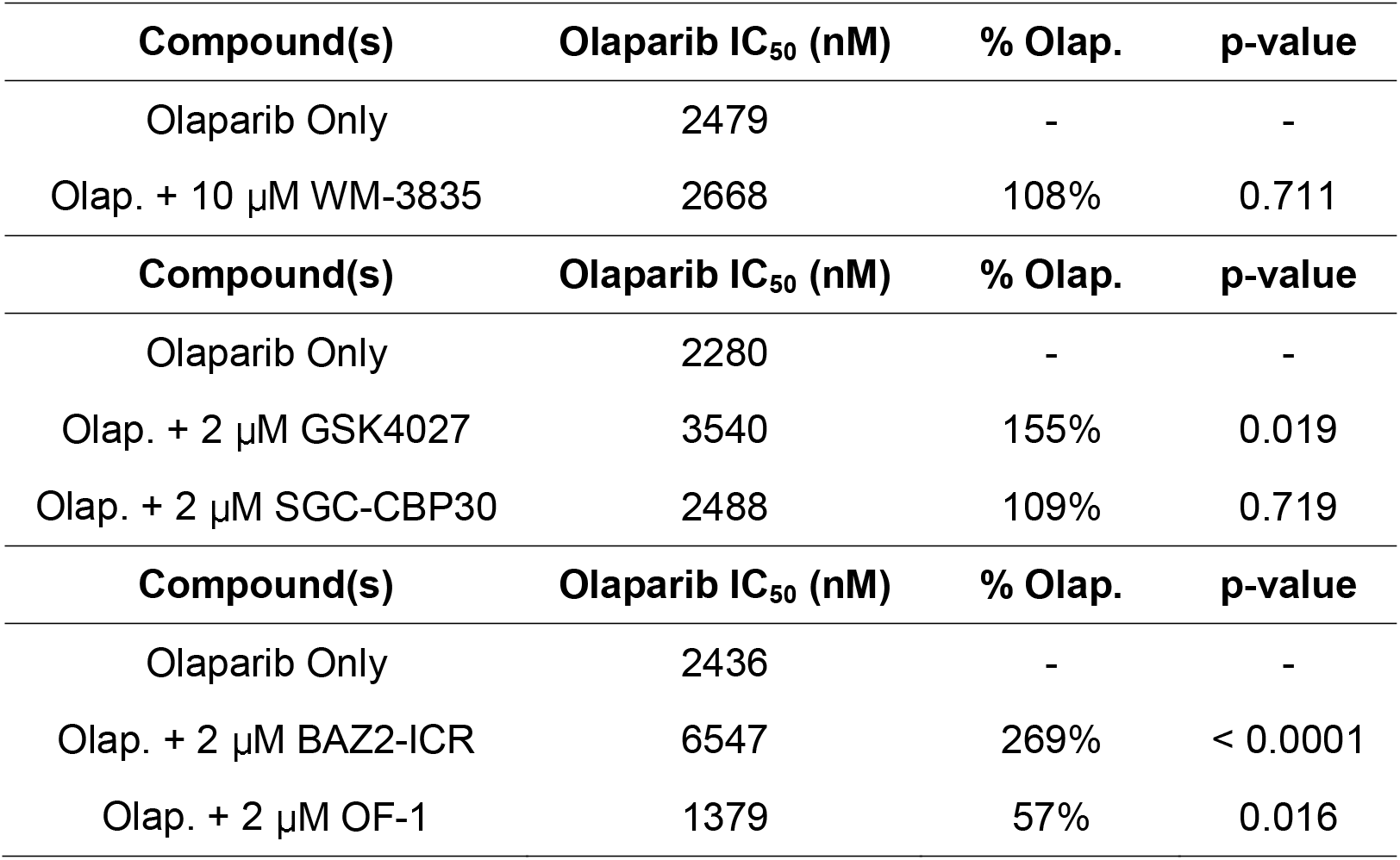
Olaparib IC_50_ values of PEO1-OR HGSOC cells treated with olaparib alone or in combination with 2 μM bromodomain inhibitors. For each treatment, the olaparib IC_50_ values are shown. Also shown are IC_50_ percentage relative to olaparib alone (% Olap.) and p-value.

| **Compound(s)** | **Olaparib IC<sub>50</sub> (nM)** | **% Olap.** | **p-value** |
| --- | --- | --- | --- |
| Olaparib Only | 2479 | - | - |
| Olap. + 10 μM WM-3835 | 2668 | 108% | 0.711 |
| **Compound(s)** | **Olaparib IC<sub>50</sub> (nM)** | **% Olap.** | **p-value** |
| Olaparib Only | 2280 | - | - |
| Olap. + 2 μM GSK4027 | 3540 | 155% | 0.019 |
| Olap. + 2 μM SGC-CBP30 | 2488 | 109% | 0.719 |
| **Compound(s)** | **Olaparib IC<sub>50</sub> (nM)** | **% Olap.** | **p-value** |
| Olaparib Only | 2436 | - | - |
| Olap. + 2 μM BAZ2-ICR | 6547 | 269% | < 0.0001 |
| Olap. + 2 μM OF-1 | 1379 | 57% | 0.016 |

While the PCAF BRD may play a role in olaparib response, it must be noted that the GCN5/PCAF BRDs also read H3K36 acetylation and the P300/CBP BRDs read H3K56 acetylation (37). We therefore chose to specifically inhibit the BRDs of proteins that have direct reading function of H3K14ac. These include Bromodomain Adjacent to Zinc Finger Domain 2B (BAZ2B) and proteins in the Bromodomain and PHD-Finger Containing (BRPF)-family (37). We performed colony formation assays in PEO1-OR cells using olaparib alone or in combination with 2 μM BAZ2B inhibitor BAZ2-ICR, or 2 μM pan-BRPF inhibitor OF-1. Much like inhibition of PCAF, inhibition of BAZ2B with BAZ2-ICR significantly increased olaparib resistance **(Fig. 3E and Table 2)**. However, inhibition of BRPF family proteins by OF-1 significantly decreased the olaparib IC_50_ **(Fig. 3F and Table 2)**. Olaparib IC_50_ and p-values for all inhibitor treatments are shown in **Table 2**.

### BRPF3 is elevated in HGSOC and associated with poor outcomes, and may play a role in olaparib response

BRPF3 is notable among the BRPF family proteins for a known association with HBO1, in which it directs a multiprotein complex toward acetylation of H3K14, rather than lysines of histone H4 (38). We examined a publicly available microarray dataset comparing mRNA expression in normal fallopian tube epithelium (FTE) versus HGSOC. *KAT7* (HBO1) mRNA levels were elevated in HGSOC **(Fig. 4A)** but results were not significant (p = 0.0833). However, *BRPF3* mRNA elevation was highly significant (p = 1.9e-06) in HGSOC compared to normal FTE **(Fig. 4B)**. We also analyzed a microarray dataset of ovarian tumors from The Cancer Genome Atlas (TCGA) and analyzed overall survival. Using KMplot (39) we split a cohort of patients into low or high expression of *KAT7* or *BRPF3*. We observed that high *KAT7* expressing patients survived longer, but the difference was not significant (p = 0.063) **(Fig. 4C)**. High *BRPF3* expressing patients, however, showed significantly shorter overall survival than low *BRPF3* expressing patients (p = 0.00056) **(Fig. 4D)**. We used the Cancer Science Institute of Singapore Ovarian Database (CSIOVDB) to perform a further meta-analysis of HGSOC microarray data (40–42). *BRPF3* was significantly upregulated in higher stage and grade of HGSOC **(Supplemental Fig. S3)**. A meta-analysis using canSAR Black (43) showed BRPF3 has the strongest association with ovarian cancer in combined molecular score, which includes mutation score, gene expression, and copy number variation score **(Supplemental Fig. S4)**.

**FIG 4.**
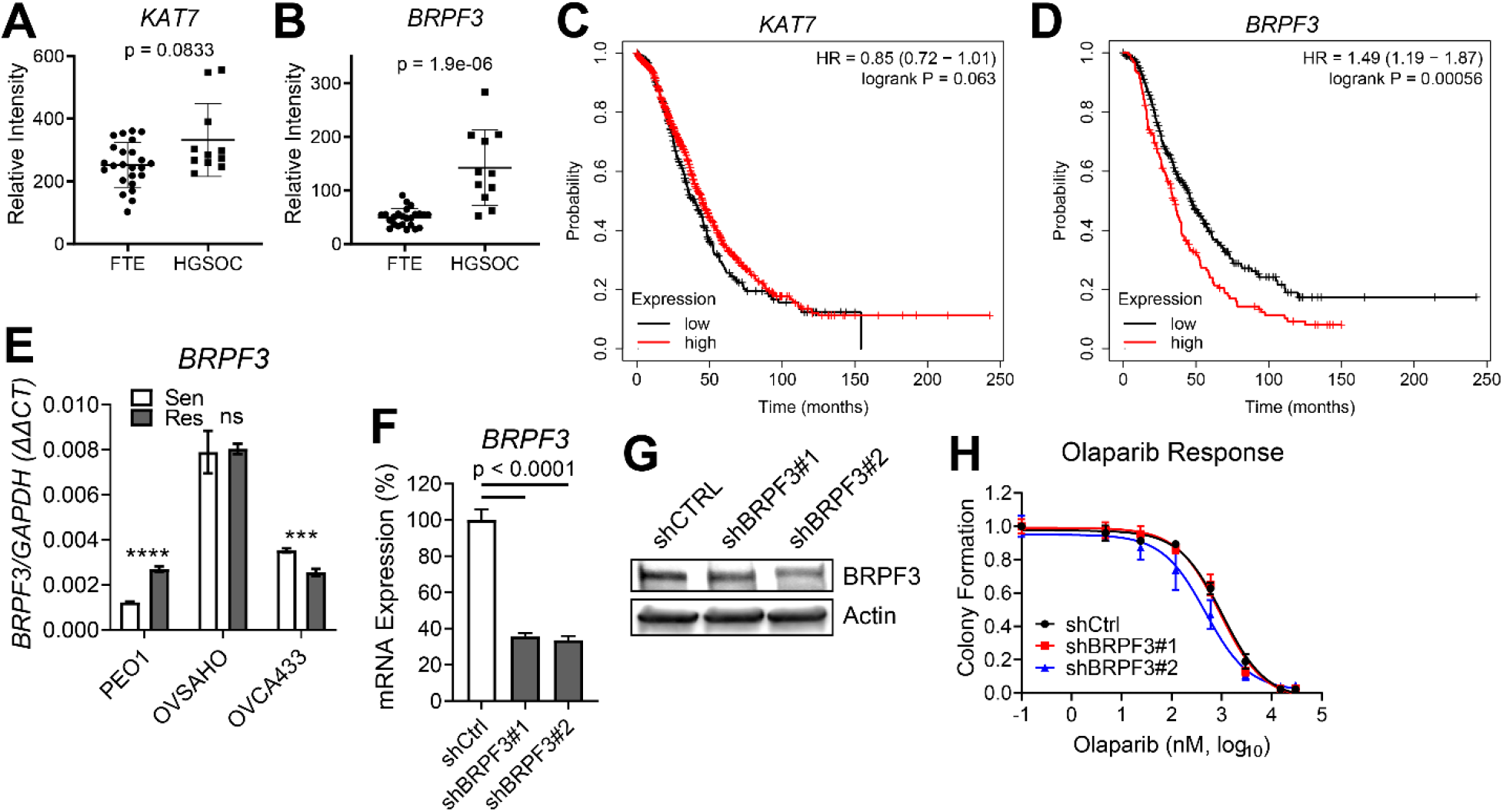
*BRPF3* mRNA expression is elevated in HGSOC relative to normal fallopian tube, and elevated expression is associated with worse overall survival. *BRPF3* knockdown moderately sensitizes PEO1-OR cells to olaparib. **(A-B)** mRNA expression of *KAT7* and *BRPF3* in microarray dataset GSE10971 was compared between HGSOC and normal FTE. p-value by t-test. **(C-D)** TCGA microarray data for *KAT7* and *BRPF3* were analyzed using KMplot and Kaplan-Meier curves were generated for overall survival (N = 655) of High and Low expressing patients for each gene. **(E)** mRNA from the indicated olaparib sensitive/resistant pairs was assayed for *BRPF3* expression by RT-qPCR. Target gene expression was determined relative to *GAPDH* control by ΔΔCt method. Each bar represents the mean ± SD of three PCR reactions. p-values by t-test. *** p < 0.001; **** p < 0.0001. **(F-G)** PEO1-OR cells were transduced with shRNA targeting *BRPF3* or a non-targeting scrambled shRNA (shCTRL) and then selected in puromycin. Knockdown of *BRPF3* was determined by RT-qPCR **(F)** and Western blotting **(G)**. mRNA expression in **(F)** was determined relative to *GAPDH* by ΔΔCt method, then normalized to shCTRL for each knockdown. p-value by t-test. **(H)** Cells from **F-G** were treated with increasing doses of olaparib, and colony formation was assayed as described in Fig. 3B.

We examined *BRPF3* mRNA expression in our three olaparib-sensitive/-resistant cell line pairs. It was increased in PEO1-OR vs sensitive PEO1, unchanged in OVSAHO-OR, and decreased in OVCA433-OR vs sensitive OVCA433 **(Fig. 4E)**. *BRPF3* was also upregulated in RNA-Seq analysis of olaparib-treated ascites of GTFB-PDX1009 **(Supplemental Table S1)**. We used lentiviral transduction to stably express shCTRL or two independent shRNA targeting *BRPF3* in PEO1-OR. We determined the degree of knockdown for each shRNA relative to shCTRL by RT-qPCR **(Fig. 4F)** and Western blot **(Fig. 4G)**. While mRNA expression appeared similar between the two *BRPF3* shRNA lines, the amount of protein in sh*BRPF3*#2 was notably lower **(Fig. 4G)**. We next compared olaparib response in *BRPF3* knockdown cells compared to control. Olaparib IC_50_ for shCTRL in this experiment was 1036 nM. sh*BRPF3*#1 (IC_50_ = 938 nM, p = 0.517) did not significantly alter olaparib IC_50_ in PEO1-OR. However, in agreement with better protein knockdown, sh*BRPF3*#2 (IC_50_ = 479 nM, p = 0.0002) did significantly reduce the olaparib IC_50_ **(Fig. 4H)**. In a duplicate experiment, the olaparib IC_50_ of shCTRL cells was 1593 nM, while the IC_50_ was significantly reduced to 835 nM in sh*BRPF3*#2 cells **(Supplemental Fig. S5)**, confirming that knockdown of *BRPF3* partially sensitized PEO1-OR to olaparib.

## Discussion

Regardless of BRCA status, numerous clinical trials showed significant progression-free survival benefit from PARPi treatment to both newly diagnosed and recurrent HGSOC cases (3, 4, 44). Going forward, nearly all HGSOC patients will be eligible to receive PARPi, and it is essential to identify and specifically target novel vulnerabilities to improve response rates and increase the number of patients who can benefit from PARPi treatment. PARPi cause ssDNA breaks and can cause replication fork collapse due to altered fork speed and stability (45, 46). Epigenetic modifications and enzymes are likely to be important mechanisms of PARPi resistance, as they control access to DNA for transcription, replication, and repair. In this report, we profiled the histone landscape of three PARPi-resistant HGSOC cell lines, including BRCA-deficient and -proficient lines. We observed increased H3K14ac in all three PARPi-resistant lines compared to isogenic PARPi-sensitive cells **(Fig. 1A-B)**. It is crucial to note that even modest changes in the percentage of a specific histone modification may represent altered enrichment or depletion at hundreds or even thousands of genomic loci in each cell and may have considerable consequences for access to DNA. Gene expression of HATs was also altered in PARPi-resistant lines, as well as in a PDX model of PARPi-treated HGSOC **(Fig. 1CD)**. Using both genetic and pharmacologic approaches, we examined if depletion or inhibition of specific HATs or accessory BRD (i.e., H3K14ac reader) proteins altered the response to the PARPi olaparib. Notably, severe pharmacologic depletion of H3K14ac did not affect PARPi response. However, knockdown or inhibition of PCAF increased PARPi resistance. Knockdown and inhibition of BRPF3, a BRD protein associated with HBO1, modestly sensitized HGSOC cells to olaparib.

Placing this work in the context of prior studies, we reported an epigenetic mechanism of PARPi resistance in HGSOC. Specifically, we previously showed that H3K9me2 was enriched in PARPi-resistant HGSOC cell line and PDX models, concomitant with upregulation of euchromatic histone-lysine N-methyltransferase-1 and −2 (EHMT1 and EHMT2), the enzymes responsible for writing the H3K9me2 mark (12). Genetic knockdown or pharmacologic inhibition of EHMT1/2 sensitized PARPi-resistant HGSOC cells and ablated both HR and NHEJ DNA repair. H3K9 methylation is known to be linked to H3K14ac by the triple Tudor domain of SETDB1, which may play a role in silencing of active genomic regions (47). We observed significant upregulation of both H3K9me2 and H3K14ac in PEO1-OR cells relative to parental, olaparib-sensitive PEO1. Profiling studies such as chromatin immunoprecipitation (ChIP) will be required to determine if regions of these two marks overlap. H3K9me2 was not as strongly upregulated in the two other olaparib-resistant lines, suggesting that a strong linkage of H3K9 methylation with H3K14ac may exist only in a subset of HGSOC. Further studies will be necessary to evaluate the efficacy of combined inhibition of EHMT1/2 with HAT or BRD inhibitors.

Knockdown of specific HATs had only modest effects at sensitizing PEO1-OR to olaparib **(Table 1)**. Despite PCAF being upregulated in PEO1-OR cells relative to olaparib-sensitive PEO1, knockdown of PCAF surprisingly further increased olaparib resistance **(Table 1 and Supplemental Fig. S2)**. In a prior study, Zhao et al. showed that acetylation of RPA1 by PCAF/GCN5 promoted NER (22), so we would expect knockdown of PCAF and/or GCN5 to lessen DNA repair and promote PARPi sensitivity. However, a separate study by Liu et al. showed that PCAF and CBP/P300 acetylate p53 in response to DNA damage (48). We may speculate that the presence of *TP53* mutations in nearly all HGSOC cases could alter this response. Additionally, Kim et al. published that PCAF acetylates H4K8 in BRCA-deficient cells at stalled replication forks and promotes fork degradation (49). Thus, knockdown of PCAF in PEO1-OR may stabilize forks and prevent cell death.

The severe depletion of H3K14ac by HBO1 inhibitor WM-3835 indicates that the majority of H3K14 acetylation in PEO1-OR cells is mediated by HBO1 **(Fig. 3A)**. A prior study by Xiao et al. demonstrated that HBO1 was enriched on the transcriptional start sites of actively transcribed genes, and that HBO1 contributed to bulk histone acetylation at these sites (50). In that study, HBO1 enrichment correlated with the level of promoter H3K14 acetylation. Given that the addition of 10 μM WM-3835 did not affect the olaparib IC_50_ in PEO1-OR **(Fig. 3B)**, we conclude that the acetyltransferase activity of HBO1, and bulk H3K14 acetylation, do not have a major effect on PARPi response. The Xiao et al. study also showed that HBO1 enrichment, despite being correlated with H3K14ac, did not correlate with transcription levels of specific genes. Thus, even though WM-3835 severely depletes H3K14ac, transcriptional data would be needed to determine if WM-3835 had significant effects on transcription. Further, despite other HATs having a lesser role in bulk genomic H3K14ac, it is possible that other HATs acetylate histones at other genomic locations and genes, some of which may alter PARPi response. We must also consider that acetylation of non-histone targets by other HATs may have effects on PARPi response. ChIP studies of specific loci and specific targeting with other HAT inhibitors will be required to tease out precise effects.

Unlike HBO1 acetyltransferase inhibition, the use of BRD inhibitors did affect olaparib response. PCAF BRD inhibitor GSK2027 increased olaparib resistance in PEO1-OR **(Fig. 3C and Table 2)**, which agrees with increased olaparib IC_50_ in PCAF knockdown cells **(Table 1)** and indicates that the BRD of PCAF, rather than the HAT domain, may play a significant role. Also, despite being readers of the same H3K14ac mark, inhibition of BAZ2B or BRPF-family proteins had opposite effects on olaparib response. Inhibition of BAZ2B with BAZ2-ICR increased resistance **(Fig. 3E and Table 2)**, while inhibition of the BRPF family with OF-1 decreased resistance **(Fig. 3F and Table 2)**, demonstrating the highly complex regulatory nature of histone modifications. Overall, we found that the blunt method of HBO1 inhibition was ineffective at reversing PARPi resistance. Changes in IC_50_ due to BRD inhibition suggest that histone readers and other accessory factors in HAT protein complexes may play more direct role in PARPi response than the HATs themselves. The exact subunit composition of such complexes in HGSOC, and how these may be altered in acquired PARPi resistance, remain unknown.

The potential importance of accessory factors is further emphasized by the increased expression of *BRPF3* in HGSOC compared to normal FTE **(Fig. 4B)**, as well as the association of elevated *BRPF3* expression with higher ovarian cancer stage and grade **(Supplemental Fig. S3)**, and with worse overall survival **(Fig. 4D)**. While OF-1 is a pan-BRPF inhibitor, specific knockdown of BRPF3 effectively reduced olaparib IC_50_ in PEO1-OR **(Fig. 4H)**, indicating its importance in PARPi response. A study by Feng et al. showed that the protein complex containing BRPF3 and HBO1 regulates H3K14 acetylation. BRPF3 was found to be crucial for H3K14 acetylation, as well as loading of CDC45 at DNA origins of replication and activation of those origins (51). Since PARPi-induced DSBs are generated during DNA replication at collapsed replication forks, this role of BRPF3 in DNA replication suggests a possible mechanism for loss or inhibition of BRPF3 to sensitize olaparib-resistant cells.

## Conclusions

Our reported findings highlight the altered epigenetic landscape of PARPi-resistant HGSOC, including histone modifications and changes in expression of associated epigenetic-modifying enzymes. While non-histone targets for acetylation have yet to be examined, our data suggest that the reader function of specific BRDs may play a more direct role than acetyltransferase domains. In conclusion, specific BRD proteins such as BRPF3, in association with HATs, may play a role in PARPi response even if total H3K14ac levels are not predictive or associated with resistance. Further examination of HAT complex composition in HGSOC and the activities of each subunit may reveal novel vulnerabilities.

## Methods

### Cell culture, shRNA, and lentivirus

Cell lines were obtained from the Gynecologic Tumor and Fluid Bank (GTFB) at the University of Colorado and were authenticated at the University of Arizona Genomics Core using short tandem repeat DNA profiling. Regular mycoplasma testing was performed using MycoLookOut PCR (Sigma). When were they last tested? HGSOC lines were cultured in RPMI 1640 supplemented with 10% fetal bovine serum (FBS) and 1% penicillin/streptomycin. 293FT lentiviral packaging cells were cultured in DMEM supplemented with 10% FBS and 1% penicillin/streptomycin. All cells were grown at 37 °C supplied with 5% CO_2_. shRNA in pLKO.1 lentiviral vector plasmid were purchased from the University of Colorado Functional Genomics Facility. Sequences and The RNAi Consortium numbers are listed in **Supplemental Table S2**. A scrambled non-targeting shRNA was used as control (Sigma-Aldrich #SHC016). Lentivirus was packaged as previously described (52) in 293FT using 3^rd^ generation packaging plasmids (Virapower, Invitrogen) with polyethyleneimene (PEI) transfection in a 1:3 DNA:PEI ratio. Culture supernatant was harvested at 48-72 hours post-transfection and processed through 0.45 μM filters. Viruses encoded a puromycin resistance gene. Transduced HGSOC cells were selected in 1 μg/mL puromycin.

### Histone modification profiling

Profiling of histone modifications in olaparib-sensitive and -resistant cells was performed by the Northwestern University Proteomics Core. Briefly, we provided frozen pellets of 5 × 10^6^ PEO1 and PEO1-OR cells. Histone extracts were trypsin digested and histone residues were assayed in triplicate as previously reported (53, 54) by liquid chromatography coupled to mass spectrometry using a TSQ Quantiva Ultra Triple Quadrupole Mass Spectrometer.

### PDX mouse model of PARPi-resistant HGSOC

All animal experiments were performed in accordance with the Guide for the Care and Use of Laboratory Animals and were approved by the University of Colorado Institutional Animal Care and Use Committee. Primary ovarian cancer sample GTFB1009 (BRCA1/2-wildtype) was provided by the University of Colorado Gynecologic Tumor and Fluid Bank. Intraperitoneal injections of 5 million GTFB1009 ascites cells were given to 6- to 8-week-old NOD SCID gamma (NSG) mice (Jackson Labs). Following a 7-day incubation period, mice were given once daily intraperitoneal injections of 50 mg/kg olaparib or vehicle control (10% 2-hydroxypropyl-β-cyclodextrin, Sigma-Aldrich #C0926) for 21 days. After treatment, tumors were allowed to recur for two months. Mice were then euthanized, and ascites were collected for analysis. This model is further characterized in (11).

### RNA-Seq of GTFB-PDX1009

RNA was isolated from control (n = 2) and olaparib-treated (n = 2) GTFB-PDX1009 ascites using the RNeasy Plus Mini kit (Qiagen). RNA quality was confirmed using an Agilent TapeStation and all RNA used for library preparation had a RIN>9. Libraries were generated and sequencing was performed by Novogene. HISAT (55) was used for alignment against GRCh37 version of the human genome. Samples were normalized using TPM (Transcripts per Kilobase Million) measurement and gene expression using the GRCh37 gene annotation was calculated using home-made scripts. Analysis was performed by the Division of Translational Bioinformatics and Cancer Systems Biology at the University of Colorado School of Medicine. Data are available at accession GSE131231.

### Compounds

P300/CBP inhibitor SGC-CBP30 (SelleckChem #S7256), PCAF inhibitor GSK4027 (Cayman #23421), and BAZ2B inhibitors GSK2801 (SelleckChem #S7231) and BAZ2-ICR (Tocris #5266) were purchased from the indicated suppliers. HBO1 inhibitor WM-3835 was kindly provided by Jonathan Baell of the Monash Institute of Pharmaceutical Sciences (Victoria, Australia) and Professor Mark Dawson of the Peter MacCallum Cancer Centre, University of Melbourne (Melbourne, Australia).

### Colony formation assay

On 24-well plates, 2500 cells were seeded and treated with increasing doses of olaparib alone or in combination with the indicated compounds. Media and drugs were changed every three days until control wells were confluent. Colonies were washed twice with PBS, then incubated in fixative (10% methanol and 10% acetic acid in PBS). Fixed colonies were stained with 0.4% crystal violet in 20% ethanol/PBS. After imaging, crystal violet was dissolved in fixative and absorbance was measured at 570 nm using a Molecular Devices SpectraMax M2e plate reader.

### Reverse-transcriptase quantitative PCR (RT-qPCR)

RNA was isolated from cells using the RNeasy Plus Mini Kit (Qiagen). mRNA expression was determined using SYBR green Luna One Step RT-qPCR Kit (New England BioLabs) on a C1000 Touch (Bio-Rad) or QuantStudio 6 (Applied Biosystems) thermocycler. Expression was quantified by the ΔΔCt method using target-specific primers and glyceraldehyde 3-phosphate dehydrogenase (*GAPDH*) control primers. mRNA-specific primers were designed to span exon-exon junctions to avoid detection of genomic DNA. Primer sequences are shown in **Supplemental Table S3**.

### Immunoblotting

For histone blots, extracts were made using the Histone Extraction Kit (Abcam #ab113476). For total protein, cells were lysed and briefly sonicated in RIPA buffer (150 mM NaCl, 1% TritonX-100, 0.5% sodium deoxycholate, 0.1% SDS, 50 mM Tris pH 8.0) supplemented with cOmplete EDTA-free protease inhibitors (Roche #11873580001) and phosphatase inhibitors NaF and Na_3_VO_4_. Protein was separated by SDS-PAGE and transferred to PVDF membrane using the TransBlot Turbo (BioRad). Membranes were blocked in LI-COR Odyssey buffer (#927-50000) for 1 hour at room temperature. Primary antibody incubation was performed overnight in blocking buffer at 4 °C. Membranes were washed 3 times for 5 minutes each in TBST (50 mM Tris pH 7.5, 150 mM NaCl, 0.1% Tween-20), then secondary antibodies were applied in blocking buffer for one hour at room temperature. Membranes were washed again 3 times for 5 minutes each in TBST. Bands were visualized using the LI-COR Odyssey Imaging System. Primary antibodies were: H3K14ac (Cell Signaling Technology Cat# 7627, RRID:AB_10839410, 1:1000), total H3 (Cell Signaling Technology Cat# 14269, RRID:AB_2756816, 1:1000), BRPF3 (Abcam Cat# ab69410, RRID:AB_1209613, 1:000), and β-actin (Abcam Cat# ab6276, RRID:AB_2223210, 1:10,000). Secondary antibodies were LI-COR fluorophore-labeled secondary goat anti-rabbit (IRDye 680RD or IRDye 800CW, LI-COR Biosciences Cat# 925-68071, RRID:AB_2721181 or LI-COR Biosciences Cat# 926-32211, RRID:AB_621843) or goat anti-mouse (IRDye 680RD or IRDye 800CW, LI-COR Biosciences Cat# 926-68070, RRID:AB_10956588 or LI-COR Biosciences Cat# 925-32210, RRID:AB_2687825) at 1:20,000 dilution.

### Densitometry

For immunoblot images captured using the LI-COR Odyssey, band fluorescence intensity was analyzed using LI-COR ImageStudio 4. Immunoblots of histone extracts were normalized to band intensity of total H3. Immunoblots of total protein lysates were normalized to intensity of β-actin.

### Ovarian cancer dataset analysis

Microarray data from publicly available ovarian cancer database GSE10971 (56, 57) was analyzed to compare gene expression between normal FTE and HGSOC. KMplot (39) was used to interrogate TCGA ovarian cancer gene expression data and compare overall survival in high and low expressing patients. CSIOVDB (40–42) and canSAR Black (43) were used to further interrogate associations between *BRPF3* and ovarian cancer microarray datasets.

### Software and statistical analysis

Statistical analysis and calculation of P value was performed using GraphPad Prism v9. Quantitative data are expressed as mean ± SD unless otherwise stated. Two-tailed t-test was used for single comparisons. Analysis of variance (ANOVA) with Fisher’s Least Significant Difference (LSD) was used in multiple comparisons. Olaparib IC_50_ were calculated and compared using the Nonlinear regression (curve fit) analysis, log (inhibitor) vs. response (three parameters) model, with extra sum-of-squares F-test. For all statistical analyses, the level of significance was set at 0.05.

## Supporting information

Complete histone mass spec data

## Declarations

### Ethics approval and consent to participate

N/A

### Consent for publication

N/A

### Availability of data and materials

Data generated and/or analyzed during this study are available from the corresponding author(s) on reasonable request. RNA-Seq dataset(s) are deposited as described in materials and methods and are publicly available.

### Author’s contributions

ZLW designed the study, performed experiments, analyzed and interpreted data, and prepared and revised the manuscript. TMY and AM generated olaparib-resistant cell lines, characterized the PDX model, and edited the manuscript. HK analyzed RNA-Seq data. BGB aided in study design, data analysis and interpretation, and manuscript preparation and revision.

## Acknowledgements

We acknowledge Professor Jonathan Baell of the Monash Institute of Pharmaceutical Sciences (Victoria, Australia) and Professor Mark Dawson of the Peter MacCallum Cancer Centre, University of Melbourne (Melbourne, Australia) and their labs for their generous gift of HBO1 inhibitor WM-3835.

## References

1. Siegel RL, Miller KD, Jemal A. Cancer statistics, 2018. CA: a cancer journal for clinicians. 2018;68(1):7–30.

2. Lisio MA, Fu L, Goyeneche A, Gao ZH, Telleria C. High-Grade Serous Ovarian Cancer: Basic Sciences, Clinical and Therapeutic Standpoints. International journal of molecular sciences. 2019;20(4).

3. Moore K, Colombo N, Scambia G, Kim BG, Oaknin A, Friedlander M, et al. Maintenance Olaparib in Patients with Newly Diagnosed Advanced Ovarian Cancer. The New England journal of medicine. 2018.

4. Domchek SM, Aghajanian C, Shapira-Frommer R, Schmutzler RK, Audeh MW, Friedlander M, et al. Efficacy and safety of olaparib monotherapy in germline BRCA1/2 mutation carriers with advanced ovarian cancer and three or more lines of prior therapy. Gynecologic oncology. 2016;140(2):199–203.

5. Matulonis UA, Penson RT, Domchek SM, Kaufman B, Shapira-Frommer R, Audeh MW, et al. Olaparib monotherapy in patients with advanced relapsed ovarian cancer and a germline BRCA1/2 mutation: a multistudy analysis of response rates and safety. Ann Oncol. 2016;27(6):1013–9.

6. Bitler BG, Watson ZL, Wheeler LJ, Behbakht K. PARP inhibitors: Clinical utility and possibilities of overcoming resistance. Gynecologic oncology. 2017;147(3):695–704.

7. Patel AG, Flatten KS, Schneider PA, Dai NT, McDonald JS, Poirier GG, et al. Enhanced killing of cancer cells by poly(ADP-ribose) polymerase inhibitors and topoisomerase I inhibitors reflects poisoning of both enzymes. The Journal of biological chemistry. 2012;287(6):4198–210.

8. Znojek P, Willmore E, Curtin NJ. Preferential potentiation of topoisomerase I poison cytotoxicity by PARP inhibition in S phase. British journal of cancer. 2014;111(7):1319–26.

9. Bajrami I, Frankum JR, Konde A, Miller RE, Rehman FL, Brough R, et al. Genome-wide profiling of genetic synthetic lethality identifies CDK12 as a novel determinant of PARP1/2 inhibitor sensitivity. Cancer research. 2014;74(1):287–97.

10. Johnson SF, Cruz C, Greifenberg AK, Dust S, Stover DG, Chi D, et al. CDK12 Inhibition Reverses De Novo and Acquired PARP Inhibitor Resistance in BRCA Wild-Type and Mutated Models of Triple-Negative Breast Cancer. Cell reports. 2016;17(9):2367–81.

11. Yamamoto TM, McMellen A, Watson ZL, Aguilera J, Ferguson R, Nurmemmedov E, et al. Activation of Wnt signaling promotes olaparib resistant ovarian cancer. Molecular carcinogenesis. 2019.

12. Watson ZL, Yamamoto TM, McMellen A, Kim H, Hughes CJ, Wheeler LJ, et al. Histone methyltransferases EHMT1 and EHMT2 (GLP/G9A) maintain PARP inhibitor resistance in high-grade serous ovarian carcinoma. Clin Epigenetics. 2019;11(1): 165.

13. Okonogi TM, Alley SC, Harwood EA, Hopkins PB, Robinson BH. Phosphate backbone neutralization increases duplex DNA flexibility: a model for protein binding. Proceedings of the National Academy of Sciences of the United States of America. 2002;99(7):4156–60.

14. Bannister AJ, Kouzarides T. Regulation of chromatin by histone modifications. Cell Res. 2011;21(3):381–95.

15. Audia JE, Campbell RM. Histone Modifications and Cancer. Cold Spring Harbor perspectives in biology. 2016;8(4):a019521.

16. Esteller M. Cancer epigenomics: DNA methylomes and histone-modification maps. Nature reviews Genetics. 2007;8(4):286–98.

17. Wang R, Xin M, Li Y, Zhang P, Zhang M. The Functions of Histone Modification Enzymes in Cancer. Curr Protein Pept Sci. 2016;17(5):438–45.

18. Lu J, He X, Zhang L, Zhang R, Li W. Acetylation in Tumor Immune Evasion Regulation. Front Pharmacol. 2021;12:771588.

19. Garcia-Ramirez M, Rocchini C, Ausio J. Modulation of chromatin folding by histone acetylation. The Journal of biological chemistry. 1995;270(30):17923–8.

20. Tse C, Sera T, Wolffe AP, Hansen JC. Disruption of higher-order folding by core histone acetylation dramatically enhances transcription of nucleosomal arrays by RNA polymerase III. Molecular and cellular biology. 1998;18(8):4629–38.

21. Duan MR, Smerdon MJ. Histone H3 lysine 14 (H3K14) acetylation facilitates DNA repair in a positioned nucleosome by stabilizing the binding of the chromatin Remodeler RSC (Remodels Structure of Chromatin). The Journal of biological chemistry. 2014;289(12):8353–63.

22. Zhao M, Geng R, Guo X, Yuan R, Zhou X, Zhong Y, et al. PCAF/GCN5-Mediated Acetylation of RPA1 Promotes Nucleotide Excision Repair. Cell reports. 2017;20(9):1997–2009.

23. Ogiwara H, Ui A, Otsuka A, Satoh H, Yokomi I, Nakajima S, et al. Histone acetylation by CBP and p300 at double-strand break sites facilitates SWI/SNF chromatin remodeling and the recruitment of non-homologous end joining factors. Oncogene. 2011;30(18):2135–46.

24. Tamburini BA, Tyler JK. Localized histone acetylation and deacetylation triggered by the homologous recombination pathway of double-strand DNA repair. Molecular and cellular biology. 2005;25(12):4903–13.

25. Ogiwara H, Kohno T. CBP and p300 histone acetyltransferases contribute to homologous recombination by transcriptionally activating the BRCA1 and RAD51 genes. PloS one. 2012;7(12):e52810.

26. Niida H, Matsunuma R, Horiguchi R, Uchida C, Nakazawa Y, Motegi A, et al. Phosphorylated HBO1 at UV irradiated sites is essential for nucleotide excision repair. Nat Commun. 2017;8:16102.

27. Matthews BG, Bowden NA, Wong-Brown MW. Epigenetic Mechanisms and Therapeutic Targets in Chemoresistant High-Grade Serous Ovarian Cancer. Cancers. 2021;13(23).

28. Sakai W, Swisher EM, Jacquemont C, Chandramohan KV, Couch FJ, Langdon SP, et al. Functional restoration of BRCA2 protein by secondary BRCA2 mutations in BRCA2-mutated ovarian carcinoma. Cancer research. 2009;69(16):6381–6.

29. Domcke S, Sinha R, Levine DA, Sander C, Schultz N. Evaluating cell lines as tumour models by comparison of genomic profiles. Nat Commun. 2013;4:2126.

30. Kuo MH, Brownell JE, Sobel RE, Ranalli TA, Cook RG, Edmondson DG, et al. Transcription-linked acetylation by Gcn5p of histones H3 and H4 at specific lysines. Nature. 1996;383(6597):269–72.

31. Grant PA, Eberharter A, John S, Cook RG, Turner BM, Workman JL. Expanded lysine acetylation specificity of Gcn5 in native complexes. The Journal of biological chemistry. 1999;274(9):5895–900.

32. Schiltz RL, Mizzen CA, Vassilev A, Cook RG, Allis CD, Nakatani Y. Overlapping but distinct patterns of histone acetylation by the human coactivators p300 and PCAF within nucleosomal substrates. The Journal of biological chemistry. 1999;274(3):1189–92.

33. Lee KK, Workman JL. Histone acetyltransferase complexes: one size doesn’t fit all. Nature reviews Molecular cell biology. 2007;8(4):284–95.

34. Nagy Z, Tora L. Distinct GCN5/PCAF-containing complexes function as co-activators and are involved in transcription factor and global histone acetylation. Oncogene. 2007;26(37):5341–57.

35. Jin Q, Yu LR, Wang L, Zhang Z, Kasper LH, Lee JE, et al. Distinct roles of GCN5/PCAF-mediated H3K9ac and CBP/p300-mediated H3K18/27ac in nuclear receptor transactivation. The EMBO journal. 2011;30(2):249–62.

36. MacPherson L, Anokye J, Yeung MM, Lam EYN, Chan YC, Weng CF, et al. HBO1 is required for the maintenance of leukaemia stem cells. Nature. 2020;577(7789):266–70.

37. Wu Q, Heidenreich D, Zhou S, Ackloo S, Kramer A, Nakka K, et al. A chemical toolbox for the study of bromodomains and epigenetic signaling. Nat Commun. 2019;10(1): 1915.

38. Feng Y, Vlassis A, Roques C, Lalonde ME, González-Aguilera C, Lambert JP, et al. BRPF3-HBO1 regulates replication origin activation and histone H3K14 acetylation. The EMBO journal. 2016;35(2):176–92.

39. Gyorffy B, Lanczky A, Szallasi Z. Implementing an online tool for genome-wide validation of survival-associated biomarkers in ovarian-cancer using microarray data from 1287 patients. Endocrine-related cancer. 2012;19(2):197–208.

40. Tan TZ, Miow QH, Huang RY, Wong MK, Ye J, Lau JA, et al. Functional genomics identifies five distinct molecular subtypes with clinical relevance and pathways for growth control in epithelial ovarian cancer. EMBO Mol Med. 2013;5(7):1051–66.

41. Tan TZ, Miow QH, Miki Y, Noda T, Mori S, Huang RY, et al. Epithelial-mesenchymal transition spectrum quantification and its efficacy in deciphering survival and drug responses of cancer patients. EMBO Mol Med. 2014;6(10):1279–93.

42. Tan TZ, Yang H, Ye J, Low J, Choolani M, Tan DS, et al. CSIOVDB: a microarray gene expression database of epithelial ovarian cancer subtype. Oncotarget. 2015;6(41):43843–52.

43. Mitsopoulos C, Di Micco P, Fernandez EV, Dolciami D, Holt E, Mica IL, et al. canSAR: update to the cancer translational research and drug discovery knowledgebase. Nucleic acids research. 2021;49(D1):D1074–d82.

44. Matulonis UA, Penson RT, Domchek SM, Kaufman B, Shapira-Frommer R, Audeh MW, et al. Olaparib monotherapy in patients with advanced relapsed ovarian cancer and a germline BRCA1/2 mutation: a multistudy analysis of response rates and safety. Ann Oncol. 2016;27(6):1013–9.

45. Maya-Mendoza A, Moudry P, Merchut-Maya JM, Lee M, Strauss R, Bartek J. High speed of fork progression induces DNA replication stress and genomic instability. Nature. 2018;559(7713):279–84.

46. D’Andrea AD. Mechanisms of PARP inhibitor sensitivity and resistance. DNA Repair (Amst). 2018;71:172–6.

47. Jurkowska RZ, Qin S, Kungulovski G, Tempel W, Liu Y, Bashtrykov P, et al. H3K14ac is linked to methylation of H3K9 by the triple Tudor domain of SETDB1. Nat Commun. 2017;8(1):2057.

48. Liu L, Scolnick DM, Trievel RC, Zhang HB, Marmorstein R, Halazonetis TD, et al. p53 sites acetylated in vitro by PCAF and p300 are acetylated in vivo in response to DNA damage. Molecular and cellular biology. 1999;19(2):1202–9.

49. Kim JJ, Lee SY, Choi JH, Woo HG, Xhemalce B, Miller KM. PCAF-Mediated Histone Acetylation Promotes Replication Fork Degradation by MRE11 and EXO1 in BRCA-Deficient Cells. Molecular cell. 2020;80(2):327–44.e8.

50. Xiao Y, Li W, Yang H, Pan L, Zhang L, Lu L, et al. HBO1 is a versatile histone acyltransferase critical for promoter histone acylations. Nucleic acids research. 2021;49(14):8037–59.

51. Feng Y, Vlassis A, Roques C, Lalonde ME, Gonzalez-Aguilera C, Lambert JP, et al. BRPF3-HBO1 regulates replication origin activation and histone H3K14 acetylation. The EMBO journal. 2016;35(2):176–92.

52. Bitler BG, Aird KM, Garipov A, Li H, Amatangelo M, Kossenkov AV, et al. Synthetic lethality by targeting EZH2 methyltransferase activity in ARID1A-mutated cancers. Nature medicine. 2015;21(3):231–8.

53. Zheng Y, Thomas PM, Kelleher NL. Measurement of acetylation turnover at distinct lysines in human histones identifies long-lived acetylation sites. Nat Commun. 2013;4:2203.

54. LaFave LM, Beguelin W, Koche R, Teater M, Spitzer B, Chramiec A, et al. Loss of BAP1 function leads to EZH2-dependent transformation. Nature medicine. 2015;21(11):1344–9.

55. Kim D, Langmead B, Salzberg SL. HISAT: a fast spliced aligner with low memory requirements. Nature methods. 2015;12(4):357–60.

56. Tone AA, Virtanen C, Shaw PA, Brown TJ. Decreased progesterone receptor isoform expression in luteal phase fallopian tube epithelium and high-grade serous carcinoma. Endocrine-related cancer. 2011;18(2):221–34.

57. Tone AA, Begley H, Sharma M, Murphy J, Rosen B, Brown TJ, et al. Gene expression profiles of luteal phase fallopian tube epithelium from BRCA mutation carriers resemble high-grade serous carcinoma. Clinical cancer research : an official journal of the American Association for Cancer Research. 2008;14(13):4067–78.

