## Supplemental Figures and Tables for "Targeting BRPF3 moderately reverses olaparib resistance in high grade serous ovarian carcinoma"

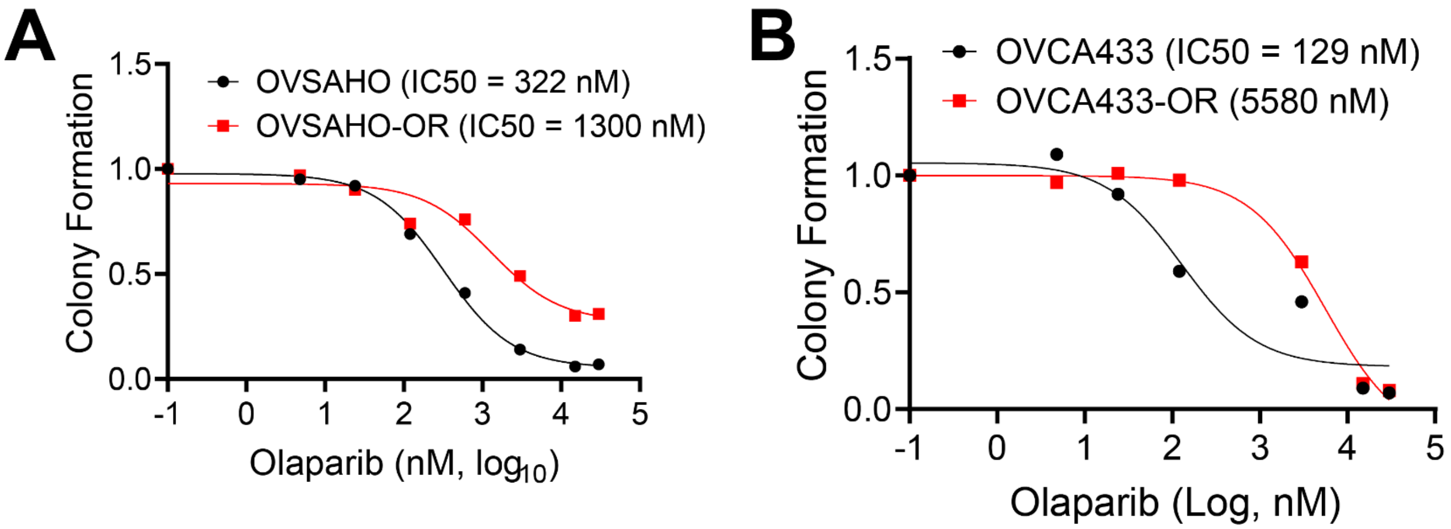


**Fig. S1. HGSOC cell lines selected in olaparib exhibit increased olaparib $IC_{50}$ relative to isogenic parental lines. (A-B)** The indicated HGSOC cell lines (OVSAHO and OVCA433) were subjected to stepwise dose escalation of olaparib to select for resistant cells. Elevated olaparib $IC_{50}$ was confirmed in olaparib-resistant (-OR) lines by dose response colony formation assays. The proportion of colonies at each dose are shown relative to vehicle control. Olaparib $IC_{50}$ values were calculated in GraphPad Prism.

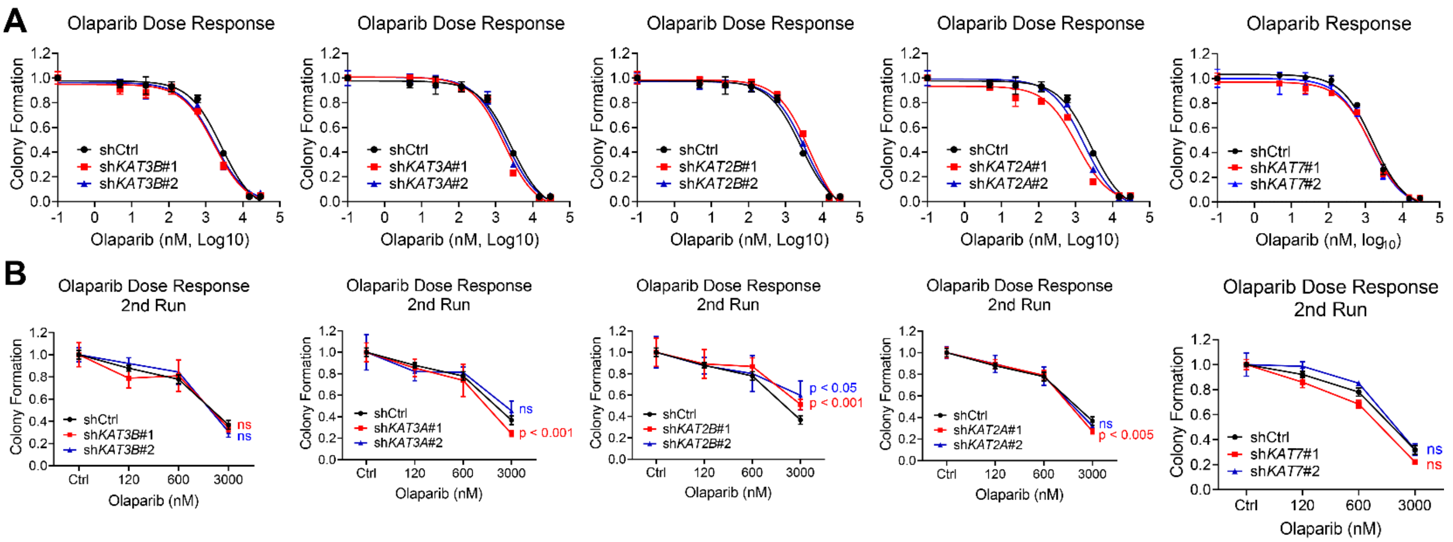


**Fig. S2. Knockdown of HATs shows only moderate alteration of olaparib response in PEO1-OR (olaparib-resistant) HGSOC cells.** PEO1-OR cells were transduced with the indicated targeted shRNAs or a non-targeting scrambled shRNA (shCTRL) and then selected in puromycin. Knockdown of each HAT was determined by RT-qPCR **(main text Fig. 2)**. **(A)** Each shRNA line was seeded at 2500 cells per well in a 24-well plate and treated with vehicle control or increasing doses of olaparib for eight days, after which colonies were fixed and stained with crystal violet. Stain was dissolved and the absorbance from each well was read by spectrophotometer, then normalized to vehicle control. $IC_{50}$ curves were plotted in GraphPad Prism. **(B)** Each shRNA line was seeded for a second dose response experiment as above, except doses were limited to vehicle control or 120 nM, 600 nM, or 3 µM olaparib. Colony formation was normalized to vehicle control for each shRNA. For all experiments, each data point is shown as mean ± SD of 3 wells. For **(B)**, p-values are calculated by t-test comparing the specific knockdown to shCtrl.

### Stage

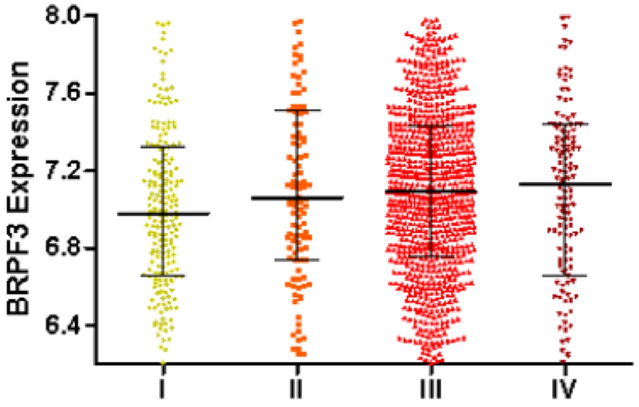


| Mann Whitney Test | I | II | III | IV | Rest |
|---|---|---|---|---|---|
| I | | 1.13e-01 | 9.70e-03 | 1.24e-01 | 1.65e-02 |
| II | | | 9.50e-01 | 7.95e-01 | 6.39e-01 |
| III | | | | 8.57e-01 | 1.40e-01 |
| IV | | | | | 8.10e-01 |

| Description\Stage | I | II | III | IV |
|---|---|---|---|---|
| Mean | 7.016 | 7.123 | 7.100 | 7.078 |
| Quantile 1 | 6.656 | 6.742 | 6.757 | 6.667 |
| Median | 6.981 | 7.062 | 7.090 | 7.128 |
| Quantile 4 | 7.324 | 7.513 | 7.436 | 7.440 |

### Grade

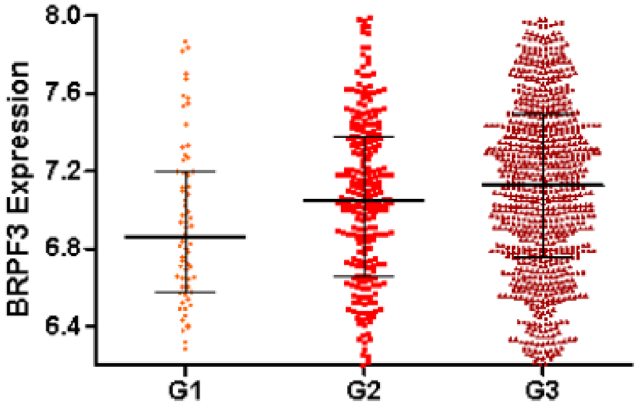


| Mann Whitney Test | Grade 1 | Grade 2 | Grade 3 | Rest |
|---|---|---|---|---|
| Grade 1 | | 5.07e-02 | 7.20e-04 | 3.76e-03 |
| Grade 2 | | | 1.13e-02 | 1.45e-01 |
| Grade 3 | | | | 5.62e-04 |

| Description\Grade | Grade 1 | Grade 2 | Grade 3 |
|---|---|---|---|
| Mean | 6.942 | 7.021 | 7.134 |
| Quantile 1 | 6.576 | 6.662 | 6.754 |
| Median | 6.861 | 7.048 | 7.130 |
| Quantile 4 | 7.196 | 7.375 | 7.493 |

**Fig. S3. BRPF3 expression is positively correlated with higher HGSOC stage and grade.** A meta-analysis of HGSOC microarray data was performed using CSIOVDB.

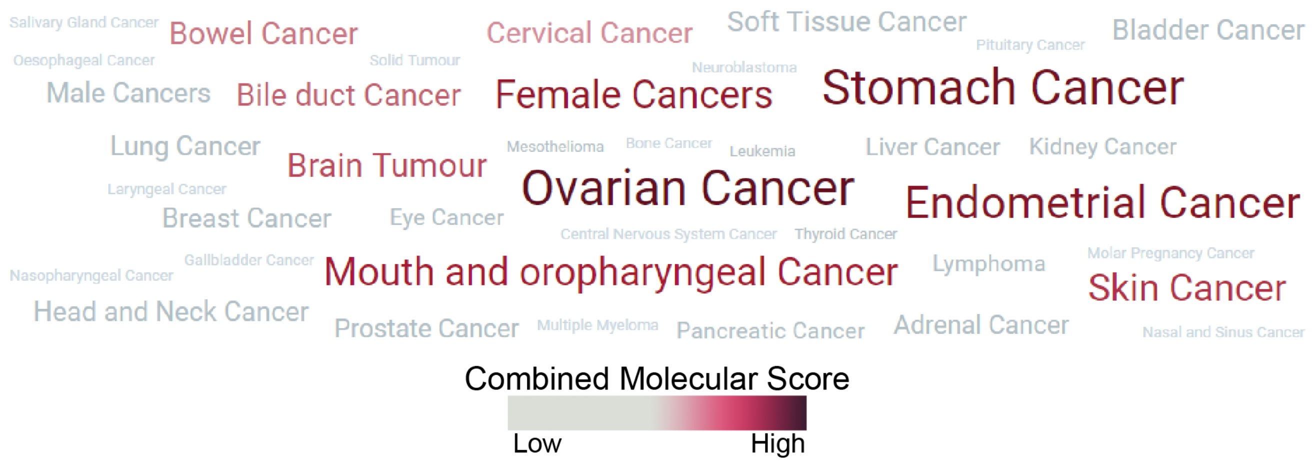


**Fig. S4. BRPF3 is significantly associated with ovarian cancer in combined molecular score.** Word cloud generated by canSAR Black by meta-analysis of cancer datasets. Combined molecular score includes mutation score, gene expression, and copy number variation.

**A** PEO1-OR Olaparib Response

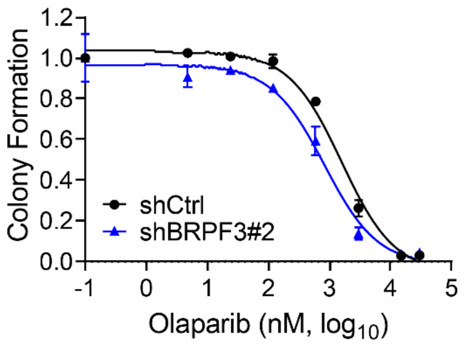


**Fig. S5. shRNA-mediated knockdown of *BRPF3* sensitizes PEO1-OR cells to olaparib.** PEO1-OR stably expressing the indicated shRNA were seeded at 2500 cells per well in a 24-well plate and treated with increasing doses of olaparib. Media and drug were changed every 2-3 days for eight days, after which colonies were fixed and stained with crystal violet. Stain was dissolved and the absorbance from each well was read by spectrophotometer, then normalized to vehicle control. Error bars, SEM. Olaparib $IC_{50}$ for each treatment condition was calculated in GraphPad Prism.

**Supplemental Tables**

| Gene (Protein) | TPM (834 Ctrl) | TPM (846 Ctrl) | TPM (827 Olap) | TPM (878 Olap) | logFC | P-value | FDR |
|---|---|---|---|---|---|---|---|
| *KAT2A* (GCN5) | 6.35 | 5.98 | 15.09 | 13.61 | 1.093 | 5.11E-05 | 0.003 |
| *KAT2B* (PCAF) | 4.92 | 3.93 | 3.63 | 4.62 | -0.149 | 0.593 | 1.000 |
| *KAT3A* (CBP) | 8.66 | 10.12 | 13.46 | 13.13 | 0.532 | 0.041 | 0.363 |
| *KAT3B* (P300) | 10.85 | 15.09 | 12.62 | 14.84 | 0.236 | 0.369 | 1.000 |
| *KAT7* (HBO1) | 7.42 | 6.56 | 6.88 | 9.45 | 0.185 | 0.497 | 1.000 |
| *BRPF3* (BRPF3) | 5 | 6.15 | 8.21 | 8.96 | 0.642 | 0.019 | 0.234 |

**Table S1. RNA-Seq analysis of PDX-GTFB1009 olaparib-treated and -untreated ascites.** For each of the indicated genes, the transcripts per million (TPM) are shown for the four indicated mice, as well as log fold change (logFC), P-value, and false discovery rate (FDR).

**Table S2. shRNA**

| **shRNA** | **The RNAi Consortium Number** | **Target Sequence** |
|---|---|---|
| shCTRL | N/A (Sigma-Aldrich #SHC016) | N/A |
| sh*KAT2A*#1 | TRCN0000038879 | GCTGAACTTTGTGCAGTACAA |
| sh*KAT2A*#2 | TRCN0000286981 | GCTGAACTTTGTGCAGTACAA |
| sh*KAT2B*#1 | TRCN0000018528 | GCAGACTTACAGCGAGTCTTT |
| sh*KAT2B*#2 | TRCN0000364134 | CCTAAACCGCATCAACTATTG |
| sh*KAT3A*#1 | TRCN0000356081 | ATCGCCACGTCCCTTAGTAAC |
| sh*KAT3A*#2 | TRCN0000356082 | CGTTTACCATGAGATCCTTAT |
| sh*KAT3B*#1 | TRCN0000231133 | TAACCAATGGTGGTGATATTA |
| sh*KAT3B*#2 | TRCN0000009883 | CAATTCCGAGACATCTTGAGA |
| sh*KAT7*#1 | TRCN0000021630 | CGGGATAAGCAGATAGAAGAA |
| sh*KAT7*#2 | TRCN0000021632 | CTCGTTCATCTGGTTCAGAAA |
| sh*BRPF3*#1 | TRCN0000382105 | AGGAGGACTTTAACCTTATAG |
| sh*BRPF3*#2 | TRCN0000277697 | TCACACTCTCTCCCGCTTATC |

**Table S3. RT-qPCR Primers**

| **Name** | **Sequence** | **Usage** |
|---|---|---|
| GAPDH_F | GTCTCCTCTGACTTCAACAGCG | Control (ΔΔCt) |
| GAPDH_R | ACCACCCTGTTGCTGTAGCCAA | Control (ΔΔCt) |
| KAT2A_GCN5_F | GCAGGTCAAGGGTTATGGGAC | Gene expression |
| KAT2A_GCN5_R | GCTCTTGGGCACCTTGATGT | Gene expression |
| KAT2B_PCAF_F | GGTGAAGAGCCATCAAAGCG | Gene expression |
| KAT2B_PCAF_R | GACTCGCTGTAAGTCTGCCA | Gene expression |
| KAT3A_CBP_F | GGCCTTCAGGTTTTGTGTGC | Gene expression |
| KAT3A_CBP_R | TTCCCAGTCTTGTGGTCTGC | Gene expression |
| KAT3B_P300_F | TGCCAAACCAGATGATGCCT | Gene expression |
| KAT3B_P300_R | ATAGCCCATAGGCGGGTTGA | Gene expression |
| KAT7_HBO1_F | CACCAACGGAGAGACAGCTT | Gene expression |
| KAT7_HBO1_R | GCAGCCTTAACTTCTCCAAATCC | Gene expression |
| BRPF3_F | CCCATCTGCAGTCCCAAAGA | Gene expression |
| BRPF3_R | GGACTTTGACCTGCTCTCGTT | Gene expression |
